# Polymer nanoparticles pass the plant interface

**DOI:** 10.1101/2022.03.24.485656

**Authors:** Sam J. Parkinson, Sireethorn Tungsirisurp, Amrita Sikder, Iseult Lynch, Rachel K. O’Reilly, Richard M Napier

## Abstract

As agriculture strives to feed an ever-increasing number of people, it must adapt to cope with climate change. It is also clear that our biosphere is suffering from an increasing burden of anthropogenic waste which includes minute plastic particles. It is not yet known whether plants will accumulate such micro- and nanoplastic materials, nor how their surface properties might influence uptake. Therefore, we prepared well-defined block copolymer nanoparticles with a range of different sizes (D_h_ = 20 - 100 nm) and surface chemistries by aqueous dispersion polymerisation using different functional macro chain transfer agents. A BODIPY fluorophore was then incorporated via hydrazone formation and uptake of these fluorescent nanoparticles into intact roots and protoplasts *of Arabidopsis thaliana* was investigated using confocal microscopy. Where uptake was seen, it was inversely proportional to nanoparticle size. Positively charged particles accumulated around root surfaces and were not taken up by roots or protoplasts, whereas negatively charged nanoparticles accumulated slowly in protoplasts and roots, becoming prominent over time in the xylem of intact roots. Neutral nanoparticles exhibited early, rapid penetration into plant roots and protoplasts, but lower xylem loads relative to the negative nanoparticles. These behaviours differ from those recorded in animal cells and our results show that, despite robust cell walls, plants are vulnerable to nanoplastic particles in the water and soil. The data form both a platform for understanding plastic waste in the farmed environment, and may also be used constructively for the design of precision delivery systems for crop protection products.

**Significance Statement:** Sustainable food production must keep pace with the growing global population, as well as adapt to climate change and other anthropogenic insults. It has become clear that micro-and nanoscale plastics are accumulating in all parts of the biosphere and we have set out to study how vulnerable plants are to such waste. We show that the size and surface properties of the designed plastics significantly affect both their speed of uptake and distribution within intact roots. Crucially, it is clear that rigid cell walls around plant cells are no barrier to the smallest particles and these pass into the plant’s vasculature. Our results relate to plastic waste but can also be used to develop precision vehicles for crop protection.

## Introduction

One of the most significant scientific advances of the 20th century was the development of synthetic polymeric materials, specifically plastics. Applications using plastics have increased exponentially, such that there is mounting concern over the contamination of marine and terrestrial environments by primary (intentionally added, e.g., as microcapsules for controlled release of agrochemicals) or secondary (breakdown products of larger plastics) micro- and nano-sized plastic particles,^1, 2^ with a Food and Agriculture Organization report emphasising the growing danger of plastics in soils.^3^ Studies on the consequences of plant exposure to nanoparticles have focused on hard particles, mostly metal-based nanoparticles plus a few dealing with structured carbon,^2, 4-8^ or chemically crosslinked polymers such as polystyrene and polyethylene beads,^9^ and so-called superabsorbent polymers (SAPs) utilised for soil remediation.^10^ However, little is known about whether or not soft, polymer-based nanoparticles will be taken up by plants, or on their transit in plants if they are accumulated. Given the increasing load on agricultural environments from the breakdown products of waste plastics and, ironically, the development of soft nano- and microplastics as precision delivery vehicles for agrochemicals,^11-14^ it is important to evaluate if these might be taken up by plants and to determine the physico-chemical properties that govern or limit uptake of these materials. Given the transition underway currently towards bio-sourced and biodegradable polymers for agricultural mulches and delivery systems, as well as plastic wastes reaching agricultural soils, model systems are needed to track and assess the implications of soft polymer nanoparticles, whose flexile structures and environmental responsiveness may allow them to be taken up and distributed in plants more effectively than their stiffer, more crystalline conterparts.^15^

Polymerisation-induced self-assembly (PISA) offers controlled soft plastic nanoparticle synthesis using block copolymers.^16^ Nanoparticle size, for example, can be managed by tuning the hydrophilic-hydrophobic block ratio. Further, PISA’s compatibility with a range of different polymerisation techniques such as reversible addition−fragmentation chain transfer (RAFT),^17, 18^ atom transfer radical polymerisation (ATRP)^19, 20^ and ring opening metathesis polymerisation (ROMP)^21, 22^ allows for a wide range of surface functionalities to be imparted to nanoparticles.

Herein, we evaluate the uptake of a series of soft plastic polymeric nanoparticles into *Arabidopsis* roots. Both intact roots and isolated, cell wall-free root protoplasts have been incubated with nanoparticles of varying small sizes (D_h_ = 20 - 100 nm) and surface functionalities (cationic, neutral and anionic). The nanoparticles were synthesised using RAFT-PISA and chemical attachment of a fluorophore allowed for their visualisation within plant tissues and cells using confocal microscopy. We demonstrate the potential of PISA for screening the impacts of systematically varying key physico-chemical characteristics (size, surface charge, shape, rigidity, hydrophobicity etc.) of soft polymeric nanoparticles as part of a safe-by-design approach for precision agriculture and greener alternatives to the current “plasticulture”.

## Results and Discussion

### Soft plastic nanoparticle synthesis and characterisation

Initially, four different macromolecular chain transfer agents (mCTAs): poly(dimethyl acrylamide) (PDMAm), poly(acrylic acid) (PAA), poly([2-(methacryloyloxy)ethyl]trimethylammonium chloride) (PQDMAEMA) and poly([2-(methacryloyloxy)ethyl]dimethyl-(3-sulfopropyl)ammonium hydroxide) (PDMAPS) were synthesised via RAFT aqueous polymerisation to allow for different surface charges to be imparted to the self-assembled polymeric nanoparticles (Figure 1, Table S1). Polymeric nanoparticles were then synthesised via RAFT aqueous dispersion polymerisation and each mCTA was chain extended with diacetone acrylamide (DAAm). For uncharged, neutral nanoparticles PDAAm degree of polymerisations (DP) of 50, 100 and 200 were targeted, the products giving hydrodynamic diameter (D_h_) values of 23 nm, 37 nm and 83 nm respectively by DLS analysis (Fig S1a) and spherical morphologies were confirmed by TEM (Figure S2).

**Figure 1.**
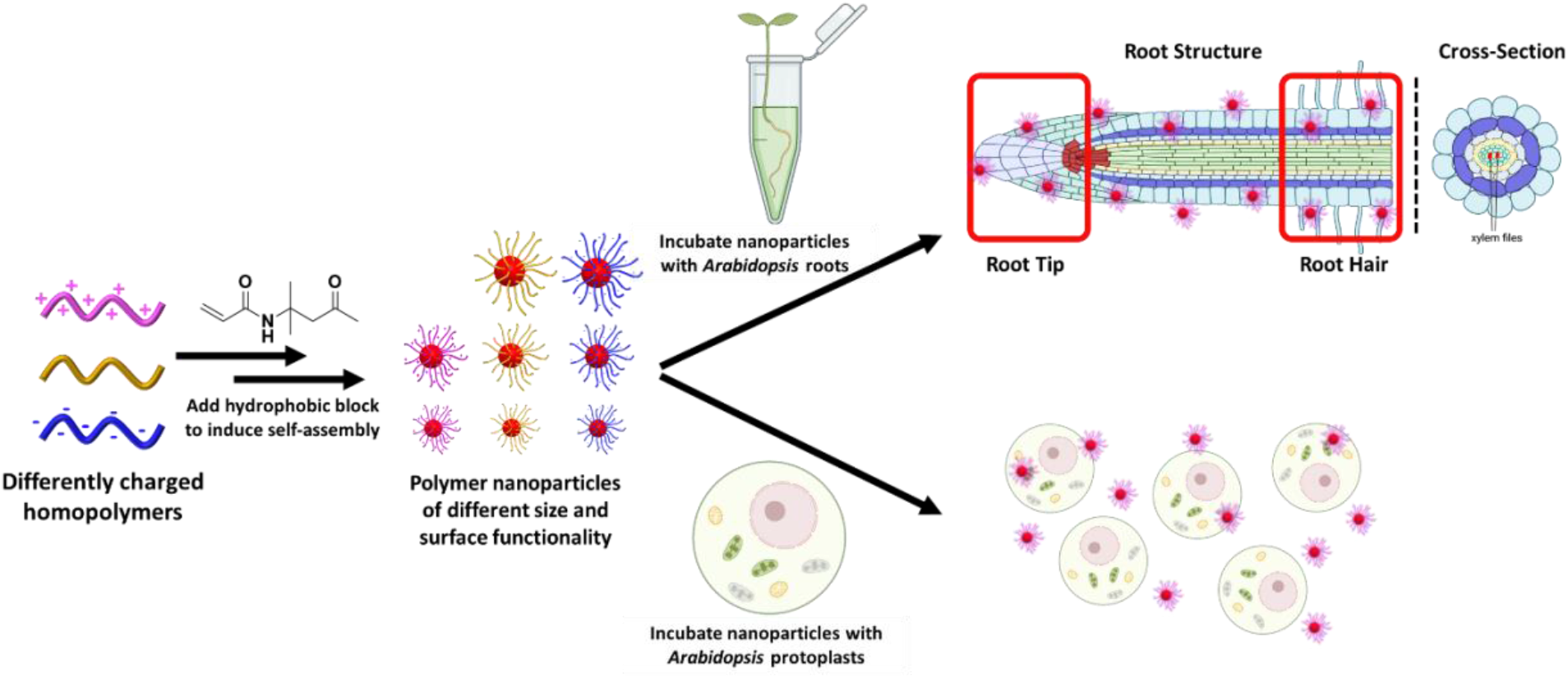
Synthesis of polymeric nanoparticles and subsequent uptake pathways explored.

During the synthesis of negative nanoparticles, it was necessary to screen the negative charges between the PAA chains by introducing PDMAm to the polymerisation reaction of DAAm. The presence of PAA and PDMAm mCTAs at a 1:9 ratio minimised the repulsion of the core acidic groups whilst maintaining an overall negative charge in the surface of these nanoparticles. The neutral PDMAm chains help to screen the negative charges between each PAA chain. With this mCTA mixture, PDAAm DPs of 50, 100 and 200 were again targeted and DLS indicated D_h_ values of 22 nm, 50 nm and 98 nm respectively (Fig S1b).

For positively charged nanoparticles, no nanoparticles were observed for PDAAm DP of 50, and it was assumed that the final polymer was still hydrophilic and that the DP was not great enough to introduce amphiphilicity to the polymer chain and, hence, induce nanoparticle assembly. For DPs 100 and 200, nanoparticles were observed with D_h_ values of 28 nm and 40 nm respectively (Fig S1c). Finally, a zwitterionic nanoparticle (D_h_ = 30 nm, Fig S1d) was synthesised for comparison to the neutral PDMAm based nanoparticles.

To visualise these nanoparticles under confocal microscopy, a fluorophore was incorporated into their core. The fluorophore needed to have an excitation wavelength away from the autofluorescence present in plant cells^23^ and BODIPY was selected because it also offers a high fluorescence quantum yield. Attachment was achieved by a hydrazone bond between the BODIPY hydrazide group and the carbonyl groups present in the PDAAm chains (Figure S3). Whilst hydrazone bonds are inherently dynamic in nature, they are stable and previous reports have shown that the use of hydrazone bonds to cross-link PDAAm chains increases particle stability.^24^

Attachment of the fluorophore was confirmed for the neutral and negative nanoparticles using size exclusion chromatography (SEC) by monitoring UV-Vis absorbance (excitation wavelength of BODIPY = 490 nm, Fig S4).

### Visualising uptake and accumulation in intact roots

In order to investigate uptake of the differently sized and charged soft nanoparticles into plants, 5-day-old *Arabidopsis thaliana* (Col-0) seedlings were incubated with each nanoparticle preparation for an hour. The accumulation (uptake and local distribution into cells) of polymer nanoparticles into the roots was monitored using confocal microscopy at both the tip, including the apex and meristematic regions (Figure S5), and further from the apex in the more mature, root hair zone (Figure 2). Plant cell walls might be considered effective physical barriers to ingress, but accumulation was seen, and penetration was found to be inversely proportional to particle size (Figure 2). Small, uncharged nanoparticles (∼20 nm) could cross the cell wall barrier and were readily taken up by the roots. Significant accumulation of neutral nanoparticles was seen inside root hairs and epidermal cells as well as at the lateral root cap and columella (Figure S5), indicating successful penetration through primary plant cell walls. These neutral nanoparticles are likely to have no electrostatic interactions with cell walls allowing them to diffuse into cells to the extent that some fluorescence was also seen in the xylem files of the vascular system (Figure 2).

**Figure 2.**
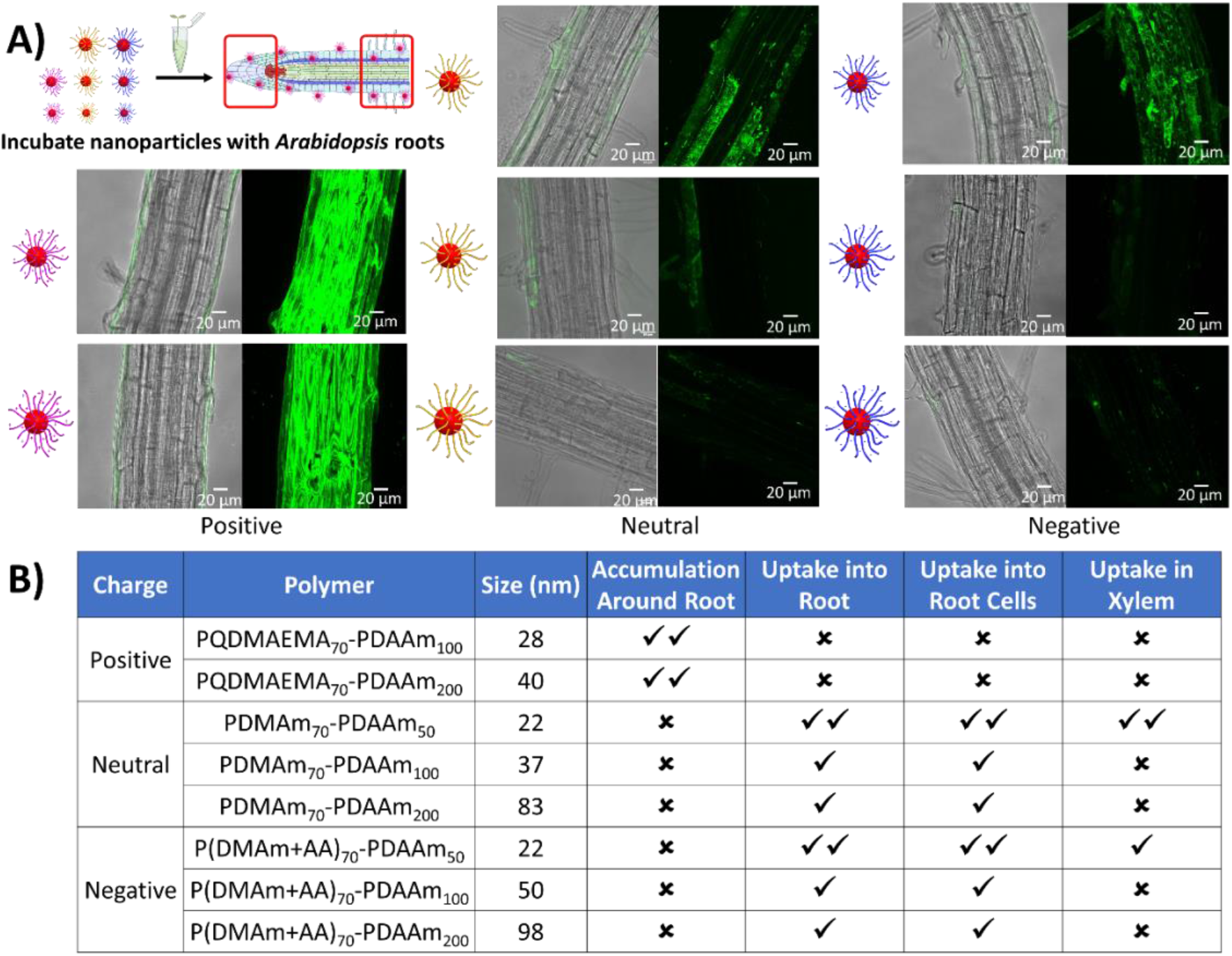
a) Confocal microscopy images for the penetration and distribution of polymeric nanoparticles in Arabidopsis root hair zones after one hour of treatment in nanoparticle solution (1 mg/ml) at room temperature. b) Summary of the different levels of root penetration observed. Good (✓✓), poor (✓) or none (✗). Penetration and accumulation were evaluated under a ZEISS 880 LSM. Maximum Z projections in 488 nm laser channel were analysed alongside the Z-slices and merged with brightfield images using ImageJ software. Scale bar = 20 μm. The images are representatives of experimental replicates (n=3).

There was a marked reduction in penetration and accumulation as the neutral nanoparticle size increased, with no fluorescence found in the xylem files for 40 nm particles or above. A moderate accumulation of fluorescence for these larger particles was detected only in the shedding lateral root cap cells and in occasional epidermal cells in the root hair zone (Figure 2). These observations were extended with our zwitterionic nanoparticles which showed similar uptake and accumulation profiles to neutral particles (Figure S6).

The smallest negatively charged nanoparticles were also accumulated into epidermal cells and root hairs (Figure 2) and accumulation was observed in shedding lateral root cap cells at the root tip (Figure S5). However, no fluorescence was detected along the xylem files in the vascular system indicating that these particles are far less mobile across and between cell layers than the neutral particles. Negatively charged nanoparticles are likely to exhibit electrostatic interactions with acidic polysaccharides and associated ions in the cell wall, affecting their accumulation and uptake.^7, 25^ As with our neutral particles, increasing particle size inhibited uptake.

Positively charged nanoparticles were observed all around the roots where they appeared to sheathe the surface. This might be due to attractive electrostatic interactions with the plant cell wall, and it appears to prohibit further passage of particles into or past the adjacent cells. This surface coating of roots has also been reported for hard nanoparticles as a function of their surface charge.^7, 25-28^. Our results are summarised in Figure 2b.

To help assess the degree of penetration of the smallest nanoparticles, orthogonal views of the confocal images were constructed. Propidium iodide (PI) staining was used to reveal the cell walls and resolve the tissue structures.^29^ These orthogonal views confirmed a sheet-like layer of positive nanoparticles (Figure 3a) coating the surface of the root, uptake of small neutral nanoparticles (Figure 3b) into both epidermal and cortical cells, and little accumulation of negative nanoparticles (Figure 3c).

**Figure 3.**
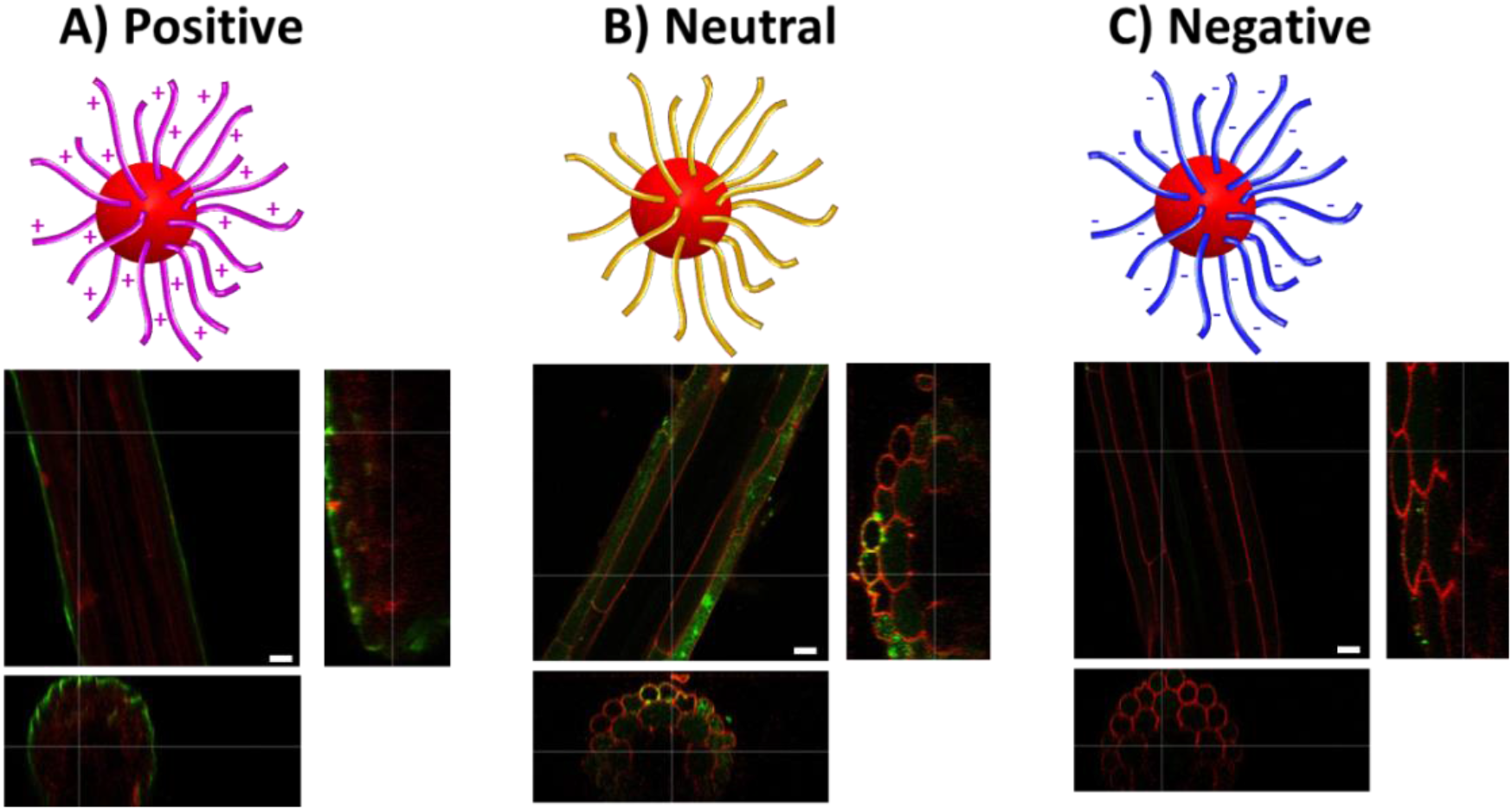
Cross-sections of root hair zones after uptake of small nanoparticles with a) positive, b) neutral and c) negative surface charges respectively. Propidium iodide staining was used to outline the cellular structure (red) while the nanoparticles are in green, and areas of co-localisation appear yellow. ImageJ software was used to generate the orthogonal views from the Z-stack images. Scale bar = 20 μm. The images are representatives of experimental replicates (n=3).

### Uptake into protoplasts

Having established that some polymer nanoparticles could penetrate into plant organs and, hence, past primary cell walls, we investigated their uptake into protoplasts (Figure 4). Protoplasts are plant cells which have had their cell walls removed enzymatically. BODIPY fluorescence emission was observed on damaged cells and cell debris regardless of the nanoparticle surface chemistry, although negative nanoparticles did show a reduced interaction with cell debris. The uptake or adsorption onto cell debris declined with increasing particle size. Interestingly, the same pattern of accumulation into the healthy (round and spherical) protoplasts was seen as in intact roots, with small neutral (Figure 4A middle) and small zwitterionic (Figure S7) nanoparticles penetrating readily. The smallest negatively charged nanoparticles accumulated in healthy protoplasts, but larger ones did not. Positively charged particles were not observed in healthy cells, but as in intact tissues they did collect around protoplasts as well as on debris. The data suggest that positive surface charge inhibits penetration through intact plant plasma membranes as well as through the cell wall.

**Figure 4.**
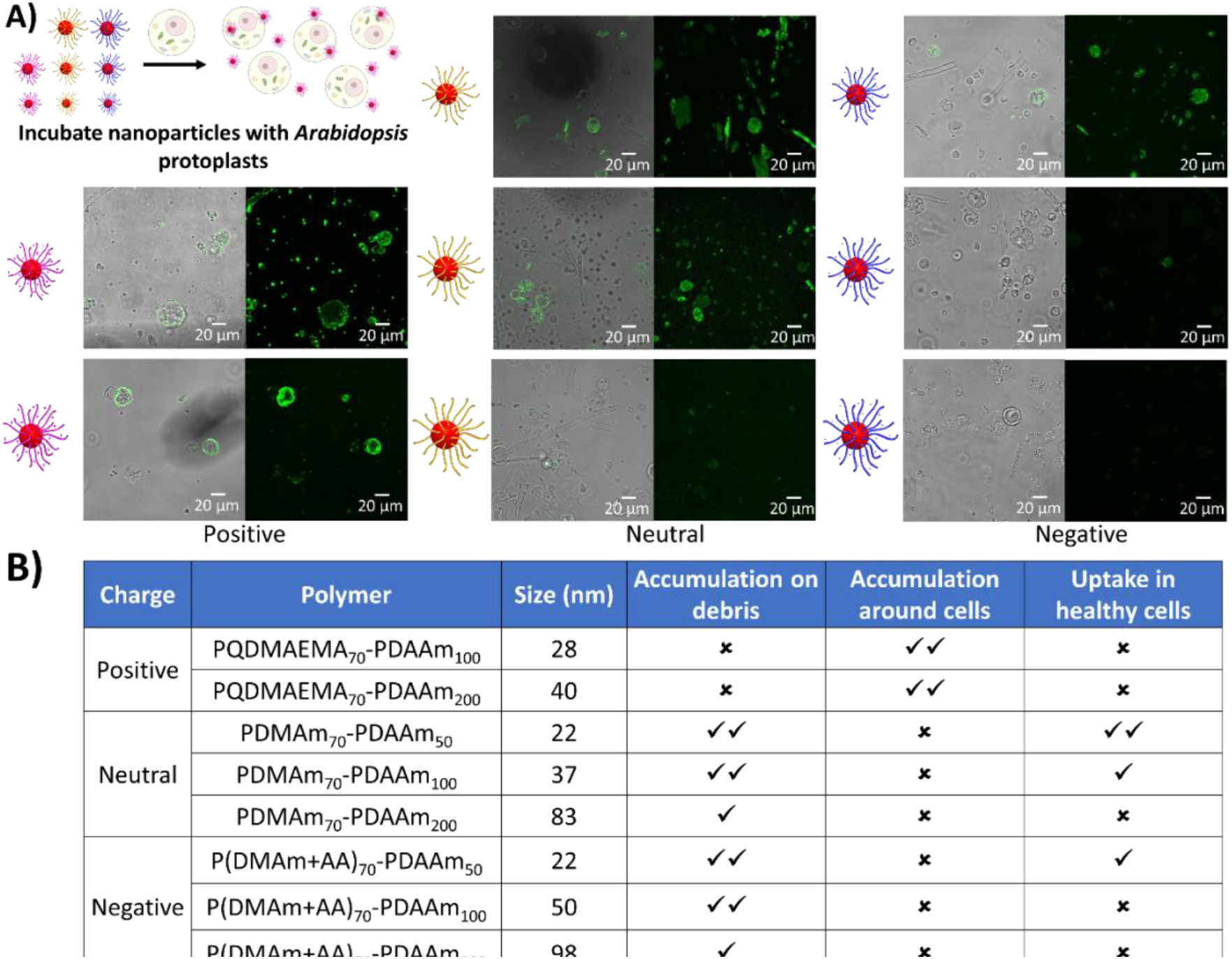
a) Confocal images for the penetration and distribution of polymeric nanoparticles by Arabidopsis protoplasts. b) Summary of the different levels of penetration observed. Good (✓✓), poor (✓) or none (✗). Penetration and accumulation were evaluated using a ZEISS 880 LSM. Maximum Z projections in the 488 nm laser channel were analysed alongside the Z-slices and merged with brightfield images using ImageJ software. Scale bar = 20 μm. The images are representatives of experimental replicates (n=3). It was noted that the smallest positively charged nanoparticles appeared to have a cytotoxic effect with an increase in visible cell debris and rough, asymmetric cells compared to other nanoparticle systems.

We may compare our data from plant exposure to soft nanoparticles to that for hard nanoparticles and to data for animal cells. In plants, gold nanoparticles, regardless of charge, were found not to pass the cell wall in *Arabidopsis*,^7^ but that both positive and negatively charged particles did enter protoplasts. Positively charged gold nanoparticles bound rapidly to the plasma membrane of tobacco protoplasts and were internalised efficiently by clathrin-mediated endocytosis, whereas negatively charged particles bound to fewer areas of membrane and were internalised by other mechanisms^30^. Further, metal oxide nanoparticles have been found to induce changes in gene expression in *Arabidopsis*.^31^ Clearly, plant cells can internalise nanoparticles if they reach the plasma membrane, and our data (Figure 4b) show that neutral particles will be internalised and accumulate rapidly.

Ready access to the plasma membrane of animal cells is similar to the case for protoplasts. As with plants, most studies have concentrated on hard particles of gold, silica, quantum dots and carbon nanotubes. Almost any nanoparticle will be taken up by animal cells, with the most effective sizes being between 40 nm and 70 nm.^32^ Uptake in this size range is dominated by clathrin-mediated endocytosis and generally positively-charged particles are internalised fastest and most effectively. Although some cell types preferentially internalise anionic particles,^33^ protein-corona binding typically results in a negative surface charge.^34^ There is some evidence that soft, small nanoparticles are internalised into cancer cells more effectively than hard particles of a similar size.^35^ Our data show clearly that there are fundamental differences between soft nanoparticle interactions with plant and animal cell systems even once the protective cell wall is breached.

### Time dependence of nanoparticle uptake

In all uptake experiments performed so far roots were incubated with nanoparticles for 1 hour and it was of interest to examine how uptake might change over time. Therefore, time series experiments were performed using the smallest nanoparticle of each functionality with incubation times ranging from 1 hour to overnight (Figure 5).

**Figure 5.**
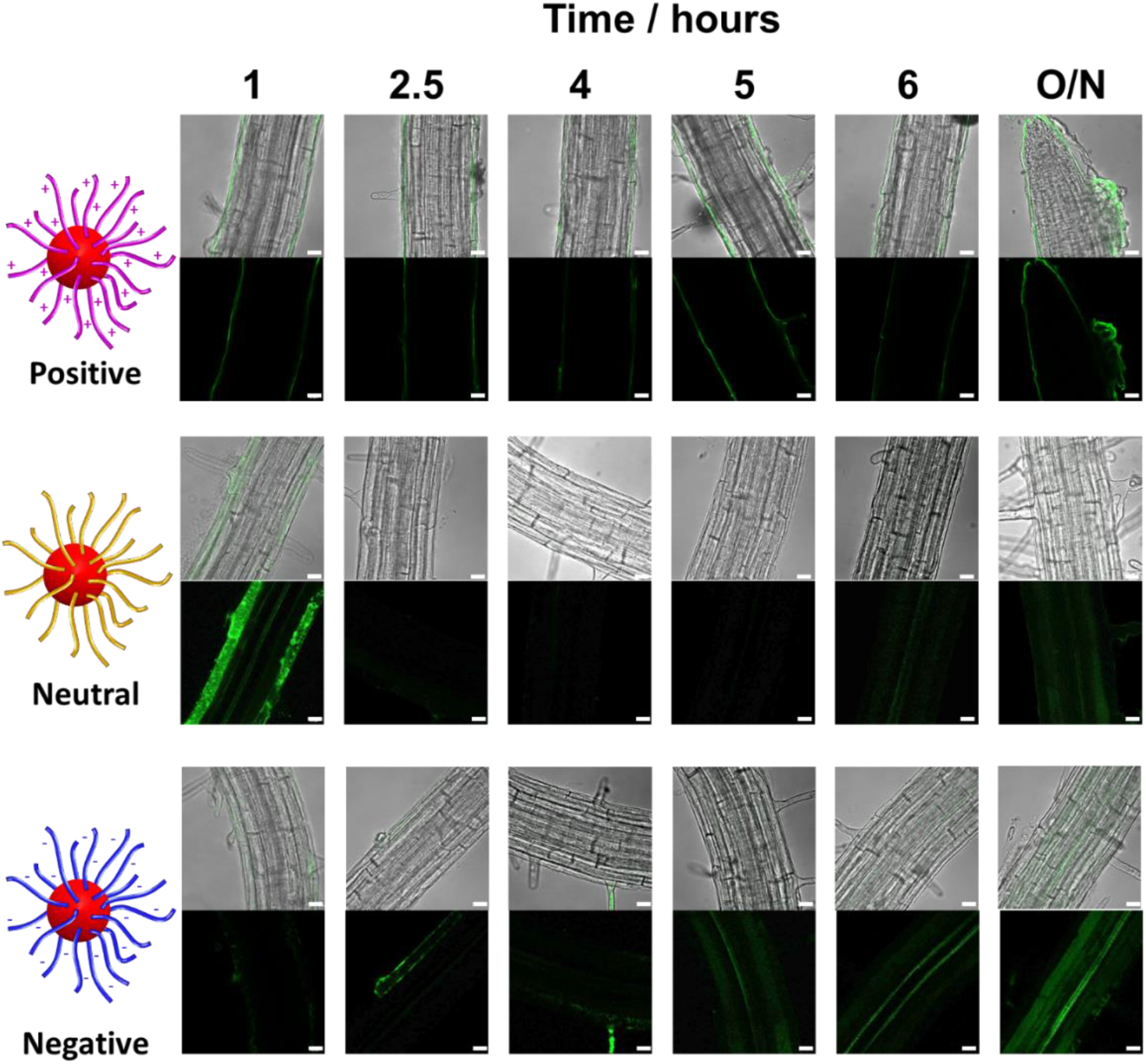
Confocal images for the penetration and accumulation of the smallest polymeric nanoparticles in Arabidopsis root hair zones over time. Penetration and accumulation were evaluated using a ZEISS 880 LSM. Maximum Z projections in the 488 nm laser channel were analysed alongside the Z-slices and merged with brightfield images using ImageJ software. Scale bar = 20 μm.

As noted before, positive nanoparticles coated the roots and this persisted through all time points (Figure 5). The accumulation of negative nanoparticles, which was low initially, was observed to increase with time. In particular, fluorescence was detected in the xylem files after 2.5 hours, becoming prominent after 5 hours and continued to increase with the overnight incubation time. These nanoparticles will thus be delivered throughout the plant in the transpiration stream. No signs of separation of the covalently-coupled BODIPY fluorophore from nanoparticles were detected when incubated in apoplastic fluid (Figure S8).

The fluorescence intensity corresponding to neutrally charged nanoparticles peaked within an hour, after which uptake appeared to decline. After 24 hours there was fluorescence in the paired xylem vessels, although the signal was weak. Similar observations were made for our zwitterionic nanoparticles without the acute early accumulation phase (Figure S9). This early burst of accumulation of nanoparticles might indicate that the plants are responding to the influx and reacting to eject or block entry as with the immune response to restrict viral distribution in plants^36^ and as seen in leaf loaded tomato plants, whereby negatively-charged nanoparticles travelling down the plant in the phloem.^37^ A low level of accumulation of neutral nanoparticles does continue giving rise to the signal in the xylem after 24 hours (Figure 5). No damage to the plants was observed.

## Concluding discussion

In summary, we have reported the uptake of polymeric nanoparticles with different sizes and surface functionalities into *Arabidopsis* root cells. By utilising RAFT-PISA, we were able to easily alter the size and functionality of these nanoparticles simply by increasing the core block DP or using a different hydrophilic macro-CTA. Attachment of a BODIPY-based fluorophore, via hydrazone bond formation with the poly(diacetone acrylamide) cores, allowed visualisation of these particles within plant tissues. Uptake into *Arabidopsis* roots and into root protoplasts was found to be inversely proportional to nanoparticle size. This differs from animal cells for which there is an optimum size range around 50 nm which equates to the mid-sized particles used here.

Given the characteristics of the cellulose matrix in plant primary cell walls the size exclusion limit reflects, in part, natural pore size limitations. However, it is notable that others have found that even very small hard nanoparticles are excluded ^7, 32^, suggesting that we need to evaluate soft polymer nanoparticles separately and with care. This seems particularly to be the case for soft particles with neutral or a negative charge. Uncharged soft plastic nanoparticles penetrated into superficial intact cells readily, and soft particles with a negative charge penetrated more slowly, but this low uptake rate persisted leading to substantial loading of the transpiration stream over time (Figure 5). It is somewhat surprising, perhaps, that once past the cell walls neutral and negatively charged nanoparticles penetrate through the plasma membrane and accumulate in the cytoplasm given the considerable turgor pressure exerted by plant cells, but our observations on protoplasts are consistent with data for gold, other metal and other hard nanoparticles and the principal mechanism appears to be clathrin-mediated endocytosis, as in animal cells.^7, 30, 38, 39^

The treatments given in this work were short, but penetration of charged polystyrene nanoparticles into plants through their roots has been reported for plants grown for a week in the treatments.^13^ By using fluorescently labelled nanoparticles (D_h_ = 200 nm), amine coated (positive) nanoparticles were found to accumulate primarily in the root epidermis of *Arabidopsis* plants, whilst sulfonate (negative) coated particles could be observed in deeper root tissues. Thus, despite their comparatively large sizes, these particles did gain access and did accumulate over long periods. Worryingly, similar but smaller (D_h_ = 55 -71 nm) particles impaired plant development.^13^ Foliar loading of small, charged star polymer nanoparticles has been studied, although most treatments included a surfactant wetting agent to combat the hydrophobicity of the plant cuticle and this is likely to have affected uptake and penetration.^37^ Once inside the tissues, both symplastic (smaller particles) and apoplastic (larger particles) transport was reported for negatively-charged polymers.

Sustainable arable agriculture must provide sufficient nutrition for the growing global population. ^40^ To date, it has generally managed to increase productivity in line with growing need. As with other parts of the biosphere, the quality of available land is decreasing due to climate change, pollution and over-fertilisation leading to low nutrient use efficiency, and so we need to ensure that we understand, as far as is possible, new and increasing threats to crop production. Nanoplastics might be a threat previously overlooked, and this work provides a foundation for continued careful evaluation of these materials. At the same time, we should recognise that nanopolymers might also provide new opportunities for reducing agrochemical inputs through precision application technologies. To this end, we must ensure that the materials developed are not damaging and that they reach their intended destinations with greatest efficiency. In this respect, if nanoparticles are to be used as vectors for crop protection, for example, the findings reported here of preferred transit to the xylem of small (<40 nm), neutral and negatively charged nanoparticles is instructive. However, the same properties potentially make these the kinds of nanoparticles most likely to pass into the food system. Armed with deeper understanding it becomes possible to tailor the application rates to optimise delivery and reduce residues, for example by further designing the soft nanoparticles to be biodegradable or hydrolysable after they have performed their useful function.

Overall, we have highlighted some of the different factors that can be used to modulate nanoparticle uptake and demonstrated RAFT-PISA as a highly versatile platform for systematic exploration of the influence of polymeric particle properties on their accumulation and localisation in plants. Subsequent work will consider other parameters such as particle stiffness (e.g., by varying the crosslink density) and impact of hydrolysable groups on retention of the nanoparticles. We believe this work begins to provide a basis for understanding how different polymeric nanoparticles will interact with plants in real-world settings.

## Supporting information

Supplemental Information incl Methods

## Acknowledgments

R.K.O, S.J.P, S.T and R.M.N acknowledge financial support from the Leverhulme Trust for funding (Grant number RPG-2016-452). A.S acknowledges funding from the European Union’s Horizon 2020 research and innovation programme under the Marie Skłodowska-Curie grant agreement no. 897666.

