## Supplemental Information incl Methods for "Polymer nanoparticles pass the plant interface"

### Experimental

#### *Materials*

Dimethyl acrylamide (DMAm, 99 %), Acrylic Acid (AA, 99 %), (Methacryloyloxy)ethyl] trimethylammonium chloride (QDMAEMA, 75 % in $H_2O$), (Methacryloyloxy)ethyl]dimethyl-(3-sulfopropyl)ammonium hydroxide (DMAPS, 95 %), 4,4'-azobis(4-cyanovaleric acid) (ACVA, 99%), 1-ethyl-3-(3-dimethylaminopropyl)carbodiimide (EDC, 98%), deuterated methanol ($CD_3OD$, 99.8%) and deuterium oxide ($D_2O$, 99.9%) were purchased from Sigma Aldrich (UK). Diacetone acrylamide (DAAm, 99%) was purchased from Alfa Aesar (UK). 4,4-Difluoro-5,7-dimethyl-4-bora-3a,4a-diaza-s-indacene-3-propionic acyl hydrazide (BODIPY FL, 99 %) was purchased from ThermoFisher Scientific (UK). 4-((((2-Carboxyethyl)thio)carbonothioyl)thio)-4-cyanopentanoic acid (BM1433, 95%) was purchased from Boron Molecular (USA).

#### *$^1$H NMR Spectroscopy*

$^1$H NMR spectra were recorded at 300 MHz on a Bruker DPX-400 spectrometer in either $D_2O$ for macro-CTA synthesis or $CD_3OD$ for all nanoparticle syntheses.

#### *Size Exclusion Chromatography (SEC)*

SEC measurements were performed on a Varian 390-LC-Multi detector suite system fitted with Refractive Index (RI) and ultraviolet (UV) detectors (λ = 309, 490 nm) equipped with a PLGel 3 µm (50 × 7.5 mm) guard column and two PLGel 5 µm (300 × 7.5 mm) mixed-D columns using DMF with 5 mM $NH_4BF_4$ at 50 °C as the eluent at a flow rate of 1.0 mL min$^{-1}$. SEC data was calibrated against polystyrene standards and analysed using Cirrus v3.3 software.

#### *Transmission Electron Microscopy (TEM)*

TEM was performed using a JEOL 2000FX or JEOL 2100FX at 200 kV. TEM solution was typically made up at 0.1 mg mL$^{-1}$ in water. Then, 10 µL of sample solution was dropped onto a carbon/formvar-coated copper grid placed on filter paper. After removing excess liquid, 10 µL of a 1% uranyl acetate solution was dropped onto the grid and left to dry.

#### *Confocal Microscopy*

*Arabidopsis* intact roots and isolated protoplasts were incubated with nanoparticle samples for at least an hour before visualisation under fluorescence confocal microscopy. For all confocal microscopy, plant samples were mounted on glass slides (1.0-1.2 mm thick) and long cover glass (22x50 mm, 0.16-0.19 mm thick) before visualisation using a Zeiss LSM 880 instrument under 40x objectives lens. The images were stacked and reconstructed by ZEN software and analysed using ImageJ software. After incubation with nanoparticles, roots were also stained with propidium iodide for 10 min and propidium fluorescence was visualised with excitation at 561 nm and emission collected from 580 – 718 nm.

#### *Synthesis of hydrophilic macro-chain transfer agents (macro-CTAs)*

A typical synthesis of a $PDMAm_{70}$ macro-CTA was as follows: dimethyl acrylamide (10 g, 100 mmol, 70 eq.), BM1433 (0.44 g, 1.4 mmol 1 eq.), ACVA (0.04 g, 140 µmol 0.1 eq.) were added to a round bottom flask and dissolved in water (24 mL) to give a 30% w/w reaction solution. A stirrer bar was added and then the flask was sealed and sparged with nitrogen for 20 minutes. The sealed flask was then immersed in an oil bath at 70 °C and left for 120 minutes after which it was removed from the oil bath and quenched by exposure to oxygen. Samples were then taken for $^1$H NMR and SEC analysis followed by purification by dialysis and then lyophilisation to yield a yellow powder. The same procedure was followed for all other macro-CTAs.

#### *Synthesis of diblock copolymer nanoparticles*

A typical synthesis of a $PDMAm_{70}$-$PDAAm_{50}$ was as follows: diacetone acrylamide (1 g, 6 mmol, 50 eq.), $PDMAm_{70}$ mCTA (0.85 g, 120 µmol 1 eq.), ACVA (3 mg, 12 µmol 0.1 eq.) were added to a round bottom flask and dissolved in water (7.4 mL) to give a 20% w/w reaction solution. A stirrer bar was added and then the flask was sealed and sparged with nitrogen for 20 minutes. The sealed flask was then immersed in an oil bath at 70 °C and left for 120 minutes after which it was removed from the oil bath and quenched by exposure to oxygen. Samples were then taken for $^1$H NMR and SEC analysis. No further purification was performed for further experiments. The same procedure was followed for all other diblock copolymer nanoparticles (see Table S1).

#### *BODIPY fluorophore attachment to polymer nanoparticles*

A typical attachment of BODIPY FL to $PDMAm_{70}$-$PDAAm_{50}$ nanoparticles was as follows: EDC (240 µg, 0.6 µmol, 1 eqv) was added to $PDMAm_{70}$-$PDAAm_{50}$ (100 mg, 0.6 µmol, 1 eq.) in water (10 mL) and stirred for 5 minutes. BODIPY FL (19 µg, 0.06 µmol, 0.1 eqv) dissolved in DMSO (15 µL) was then added and the solution was left to stir overnight. The solution was then purified by spin centrifugation against a 3k MWCO membrane. A sample was taken for SEC to confirm attachment of BODIPY to the polymer chains. The same procedure was followed for all other nanoparticles.

#### *Arabidopsis thaliana root preparation*

*Arabidopsis thaliana* growth conditions: *A. thaliana* ecotype Columbia-0 was used for all nanoparticle uptake experiments. The seeds were surface-sterilised using 10% Bleach and 70% ethanol prior to seeding on sterile ½ strength Murashige and Skoog (MS) medium (2.2 g/L MS supplemented with B5 vitamins, 1% sucrose, 1% agar, pH 5.8). Seeds were stratified on ½ MS media agar plate in the dark for two days at 4°C prior to germination for 5 days at 22°C with 12-hour daytime.

#### *Protoplast isolation*

Protoplast solution (600 mM Mannitol, 2 mM $MgCl_2$, 2 mM $CaCl_2$, 10 mM KCl, 2 mM MES, 0.1% w/v bovine serum albumin) was prepared and adjusted to pH 5.5 with Tris-HCl, 0.2 µm filtered, and stored at -20°C until use. To isolate protoplasts, 5-day-old *Arabidopsis thaliana* roots were transferred into enzyme solution (1.5% w/v Cellulase RS, 0.1% w/v Pectolyase in protoplast solution) and chopped finely using a sterile blade. The solution with chopped roots was transferred to a 35 mm round Petri dish and incubated in the dark at 25°C with constant agitation for at least 2 hr. The resulting cell suspension was filtered sequentially through 70 µm and 40 µm meshes pre-soaked with enzyme solution. The protoplasts in solution were carefully transferred to polystyrene culture tubes and an equal volume of fresh protoplast solution added, before centrifugation at 300 g for 5 min at 4°C. The supernatant was discarded, the protoplasts resuspended in the same volume of protoplast solution and centrifugation repeated. The protoplasts were resuspended in 500 µL protoplast solutiona and used directly.

**Table S*1*.** Characterisation data obtained for polymer nanoparticles incubated with *Arabidopsis thaliana* roots

| *Polymer* | *$M_n$ / g mol$^{-1}$* [a] | *Đ (GPC)* | *$D_h$ / nm* [b] | *Đ (DLS)* |
|---|---|---|---|---|
| $PDMAm_{70}$-$PDAAm_{50}$ | 36,500 | 1.14 | 23 | 0.06 |
| $PDMAm_{70}$-$PDAAm_{100}$ | 48,200 | 1.14 | 37 | 0.02 |
| $PDMAm_{70}$-$PDAAm_{200}$ | 61,500 | 1.15 | 83 | 0.08 |
| $P(DMAm_{70}+AA_{70})$-$PDAAm_{50}$ | 36,800 | 1.17 | 22 | 0.14 |
| $P(DMAm_{70}+AA_{70})$-$PDAAm_{100}$ | 51,100 | 1.15 | 50 | 0.09 |
| $P(DMAm_{70}+AA_{70})$-$PDAAm_{200}$ | 57,500 | 1.22 | 98 | 0.07 |
| $PQDMAEMA_{70}$-$PDAAm_{100}$ | N/A | N/A | 28 | 0.13 |
| $PQDMAEMA_{70}$-$PDAAm_{200}$ | N/A | N/A | 40 | 0.09 |
| $PDMAPS_{70}$-$PDAAm_{50}$ | N/A | N/A | 30 | 0.08 |

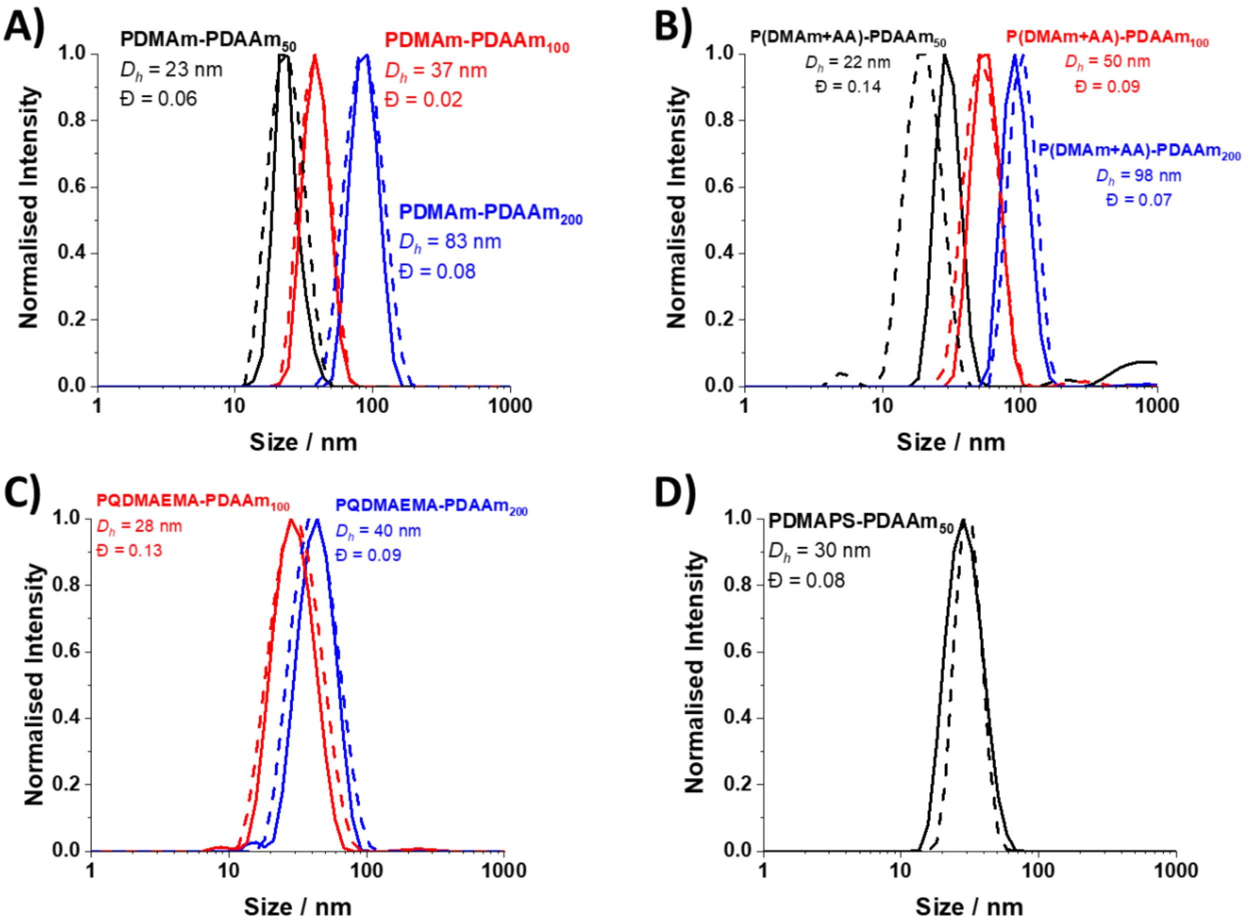


**Figure S1.** DLS traces for a) PDMAm-PDAAm, b) P(DMAm+AA)-PDAAm, c) PQDMAEMA-PDAAm and d) PDMAPS-PDAAm nanoparticles both before (dashed) and after (solid) BODIPY attachment.

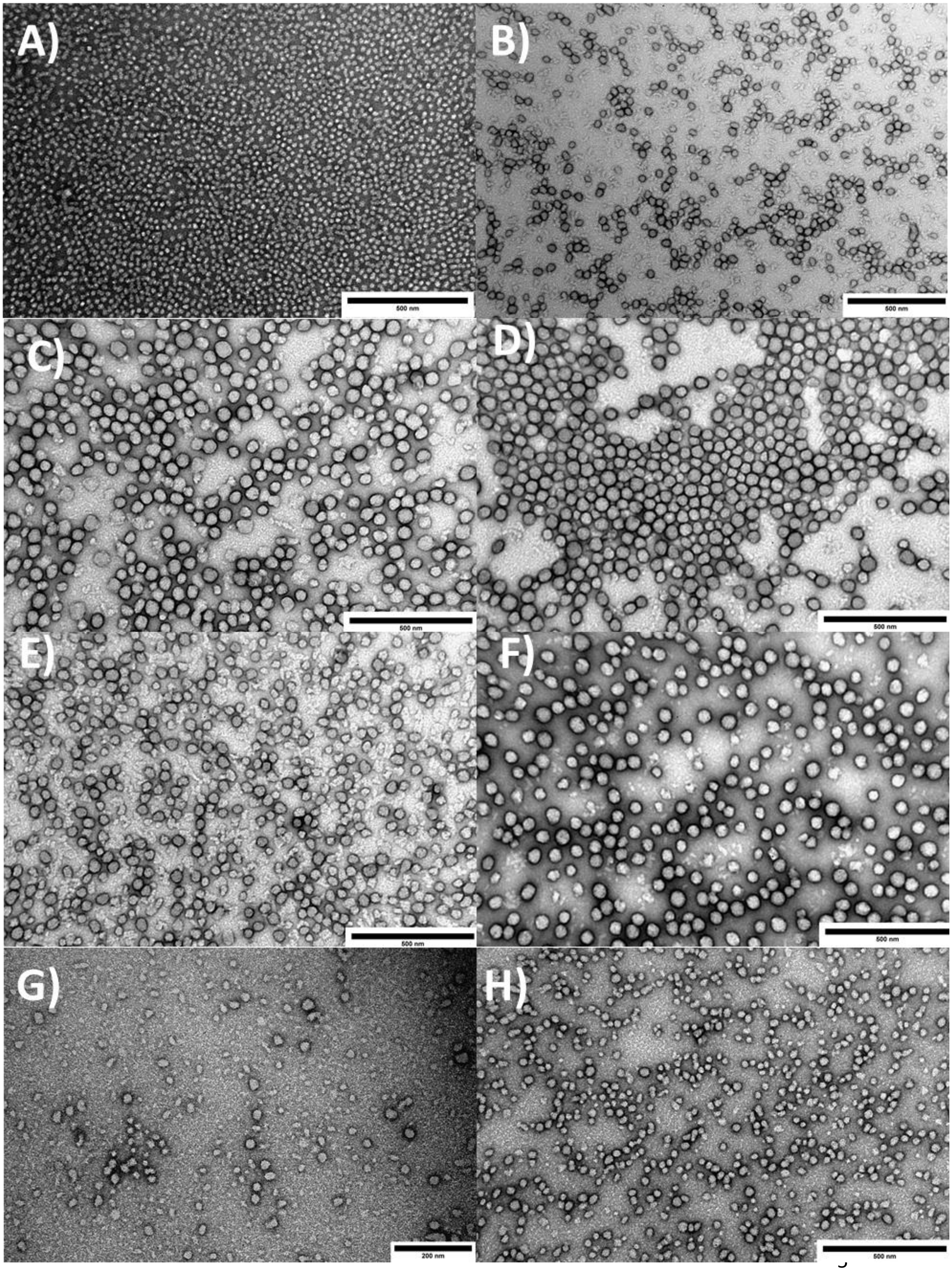
A)
500 nm
B)
500 nm
C)
500 nm
D)
500 nm
E)
500 nm
F)
500 nm
G)
200 nm
H)
500 nm

**Figure S2.** TEM images for a) PDMAm-PDAAm$_{50}$, b) PDMAm-PDAAm$_{100}$, c) PDMAm-PDAAm$_{200}$, d) P(DMAm+AA)-PDAAm$_{50}$, e) P(DMAm+AA)-PDAAm$_{100}$, f) P(DMAm+AA)-PDAAm$_{200}$, g) PQDMAEMA-PDAAm$_{100}$, h) PQDMAEMA-PDAAm$_{200}$.

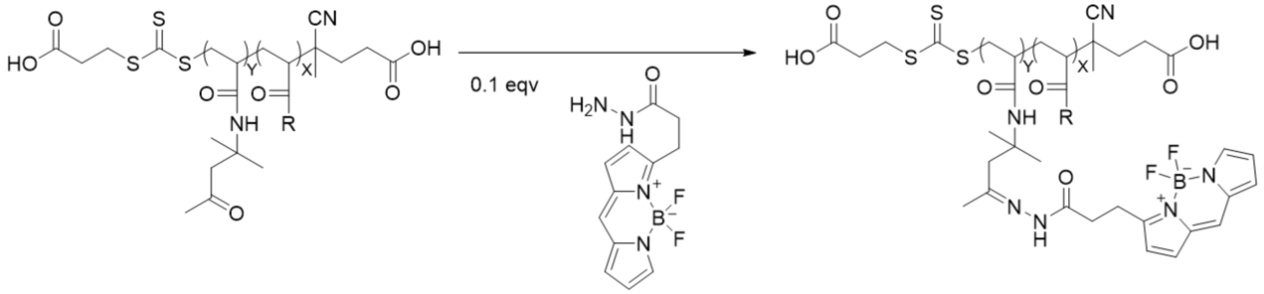


**Figure S3.** Reaction scheme for BODIPY fluorophore attachment to polymeric nanoparticles.

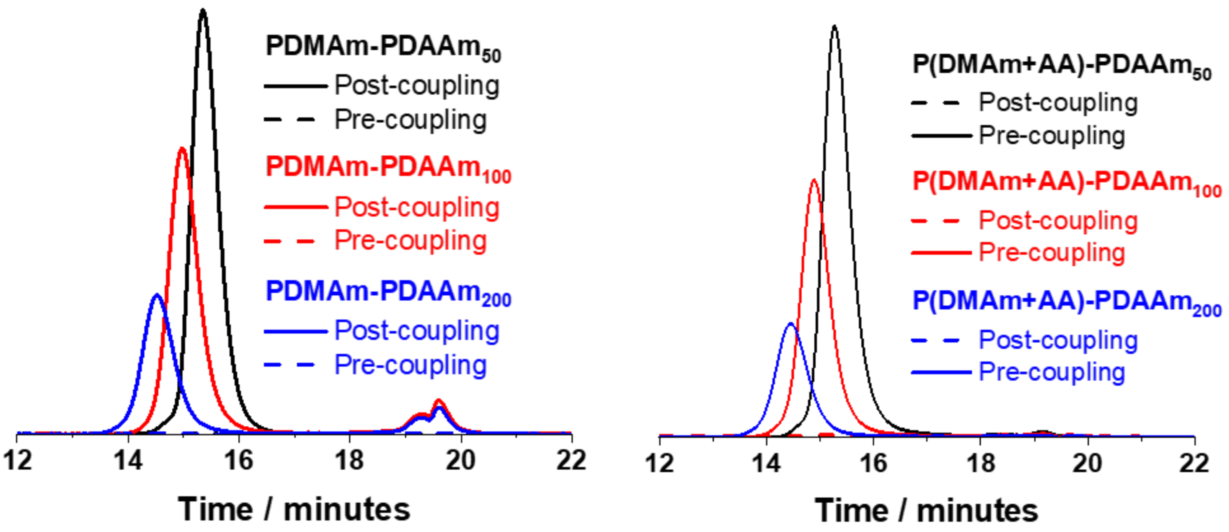


**Figure S4.** UV-SEC traces obtained at 490 nm for a) PDMAm-PDAAm and b) P(DMAm+AA)-PDAAm nanoparticles before (dashed) and after (solid) BODIPY fluorophore attachment.

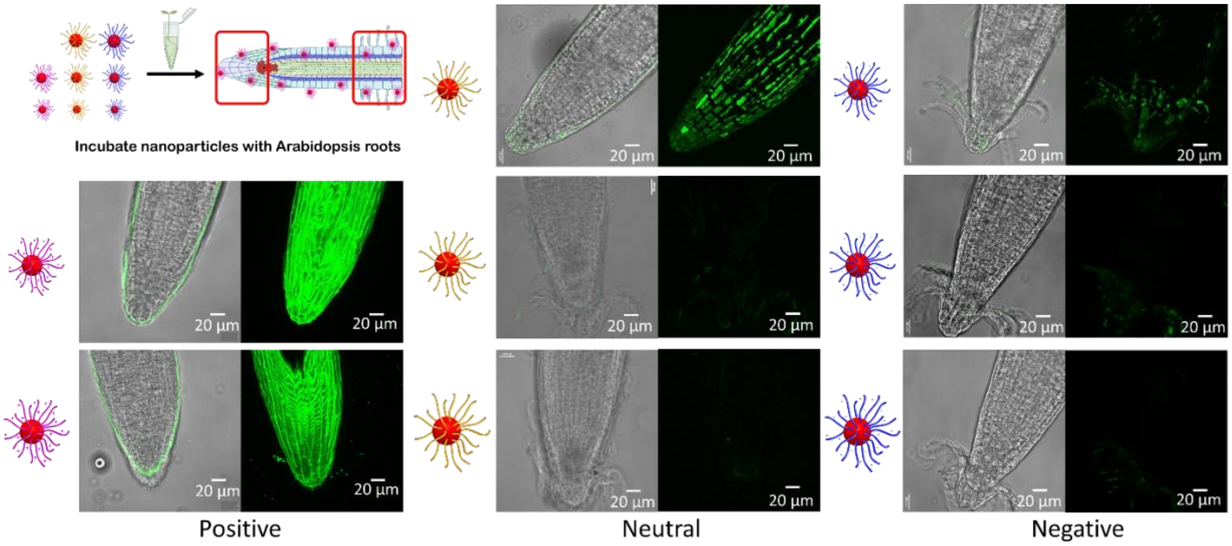


**Figure S5.** Confocal images for the penetration (entry into cells) and accumulation (extent of nanoparticle build up in cells) of polymeric nanoparticles by *Arabidopsis* root tips. Penetration and accumulation were evaluated using a ZEISS 880 LSM. Maximum Z projections in the 488 nm laser channel were analysed alongside the Z-slices and merged with brightfield images using ImageJ software. Scale bar = 20 µm. The images are representatives of experimental replicates (n=3).

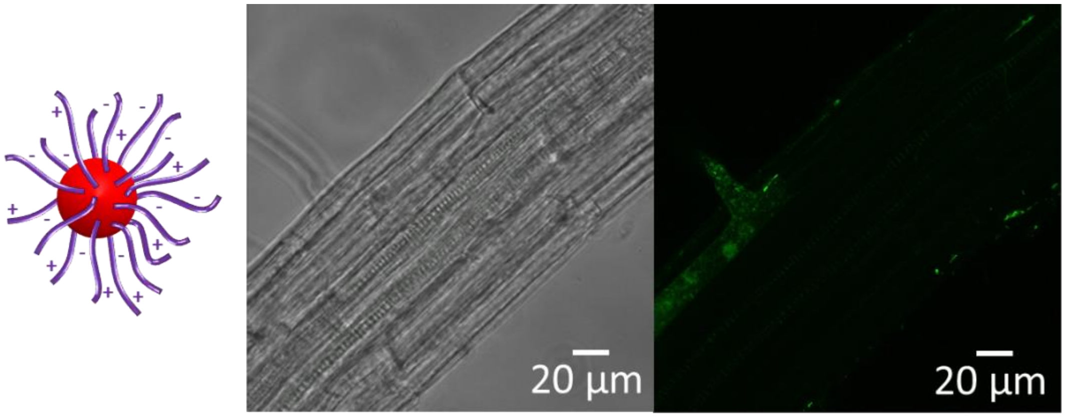


**Figure S6.** Confocal images for the penetration and accumulation of zwitterionic polymeric nanoparticles in *Arabidopsis* root hair zones. Penetration and accumulation were evaluated under ZEISS 880 LSM. Maximum Z projections in 488 nm laser channel were analysed alongside the Z-slices and merged with brightfield images using ImageJ software. Scale bar = 20 µm. The images are representatives of experimental replicates (n=3).

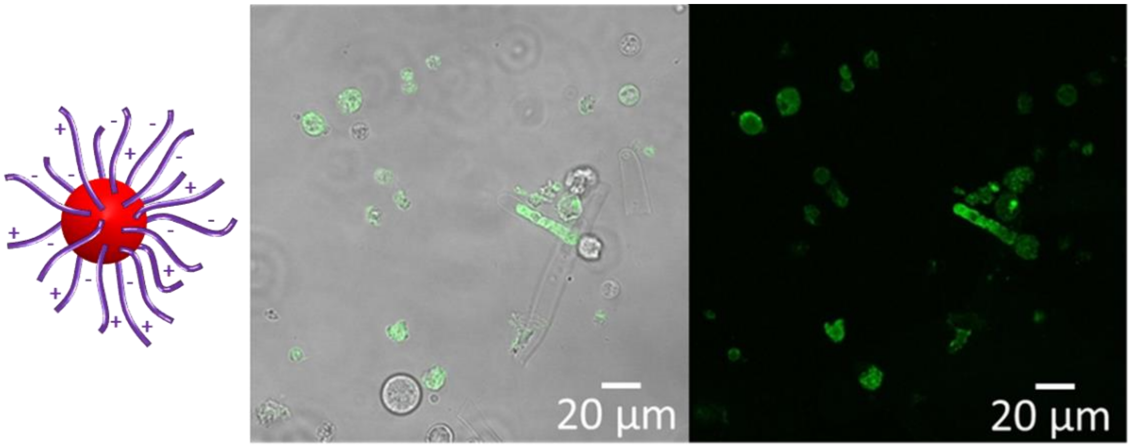


**Figure S7.** Confocal images for the penetration and accumulation of zwitterionic polymeric nanoparticles in *Arabidopsis* protoplasts. Penetration and accumulation were evaluated under ZEISS 880 LSM. Maximum Z projections in 488 nm laser channel were analysed alongside the Z-slices and merged with brightfield images using ImageJ software. Scale bar = 20 µm. The images are representatives of experimental replicates (n=3).

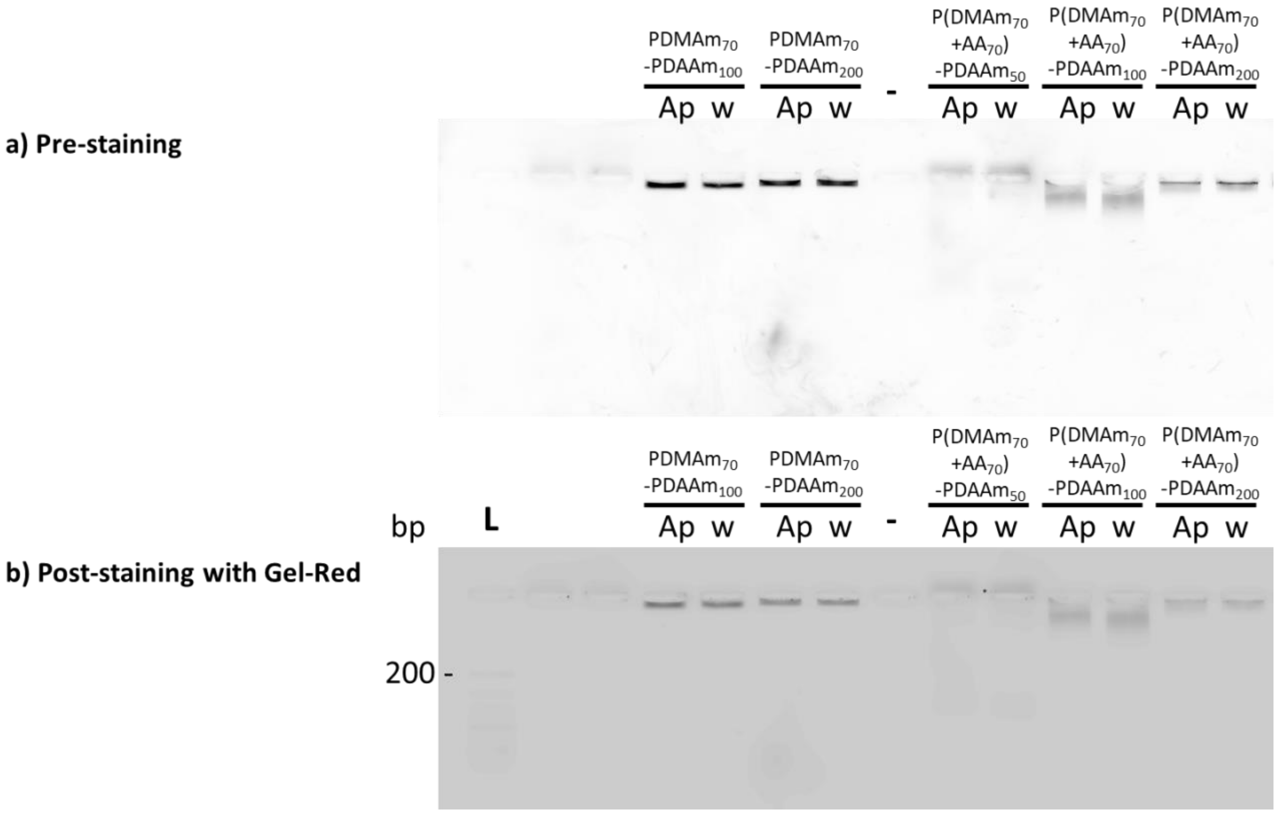

**Figure S8.** Comparable stability of CPNs upon incubation in extracted apoplastic fluid from *Nicotiana benthamiana* leaves (Ap) in comparison to those incubated in distilled water (w). 2% agarose gel electrophoresis was performed with low $M_w$ DNA ladder (L) as size standard. A negative control of apoplastic fluid alone (-) confirmed no background bands. Incubations were for 2.5 hours at room temperature. The gel was imaged using a Typhoon fluorescence imager with filters for BODIPY (top panel) before it was stained with Gel-Red and imaged again using the RGB filter. No bleeding of the BODIPY away from the nanoparticle bands was observed.

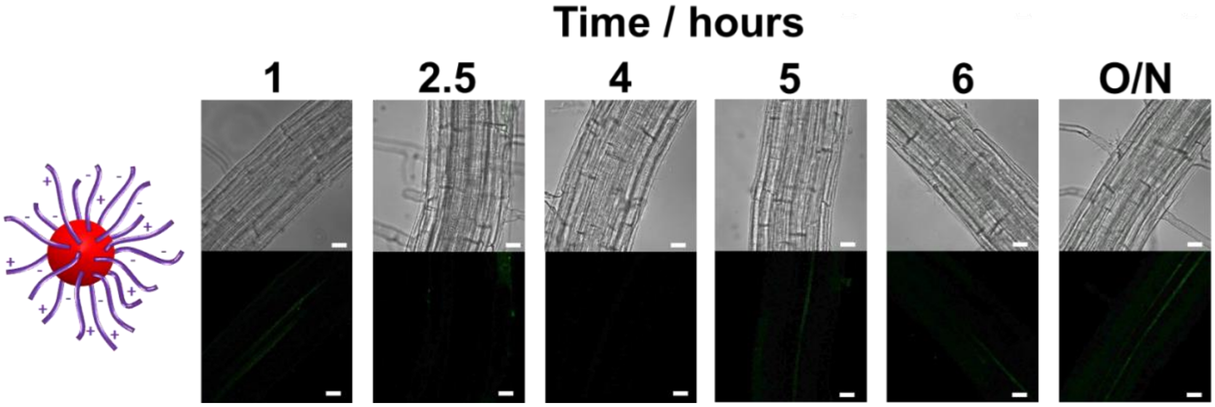


**Figure S9.** Confocal images for the penetration and accumulation of zwitterionic polymeric nanoparticles in *Arabidopsis* root hair zones over time. Penetration and accumulation were evaluated under ZEISS 880 LSM. Maximum Z projections in 488 nm laser channel were analysed alongside the Z-slices and merged with brightfield images using ImageJ software. Scale bar = 20 µm. The images are representatives of experimental replicates (n=3).
